# Computational staining of pathology images to study tumor microenvironment in lung cancer

**DOI:** 10.1101/630749

**Authors:** Shidan Wang, Ruichen Rong, Donghan M. Yang, Ling Cai, Lin Yang, Danni Luo, Bo Yao, Lin Xu, Tao Wang, Xiaowei Zhan, Yang Xie, Adi Gazdar, John Minna, Guanghua Xiao

## Abstract

The spatial organization of different types of cells in tumor tissues reveals important information about the tumor microenvironment (TME). In order to facilitate the study of cellular spatial organization and interactions, we developed a comprehensive nuclei segmentation and classification tool to characterize the TME from standard Hematoxylin and Eosin (H&E)-stained pathology images. This tool can computationally “stain” different types of cell nuclei in H&E pathology images to facilitate pathologists in analyzing the TME.

A Mask Regional-Convolutional Neural Network (Mask-RCNN) model was developed to segment the nuclei of tumor, stromal, lymphocyte, macrophage, karyorrhexis and red blood cells in lung adenocarcinoma (ADC). Using this tool, we identified and classified cell nuclei and extracted 48 cell spatial organization-related features that characterize the TME. Using these features, we developed a prognostic model from the National Lung Screening Trial dataset, and independently validated the model in The Cancer Genome Atlas (TCGA) lung ADC dataset, in which the predicted high-risk group showed significantly worse survival than the low-risk group (pv= 0.001), with a hazard ratio of 2.23 [1.37-3.65] after adjusting for clinical variables. Furthermore, the image-derived TME features were significantly correlated with the gene expression of biological pathways. For example, transcription activation of both the T-cell receptor (TCR) and Programmed cell death protein 1 (PD1) pathways was positively correlated with the density of detected lymphocytes in tumor tissues, while expression of the extracellular matrix organization pathway was positively correlated with the density of stromal cells.

This study developed a deep learning-based analysis tool to dissect the TME from tumor tissue images. Using this tool, we demonstrated that the spatial organization of different cell types is predictive of patient survival and associated with the gene expression of biological pathways. Although developed from the pathology images of lung ADC, this model can be adapted into other types of cancers.

## 1 INTRODUCTION

With the advance of technology, Hematoxylin and Eosin (H&E)-stained tissue slide scanning has become a routine clinical procedure, which produces pathology images that capture histological details in high resolution. Pathology images of tumor tissues contain not only essential information for tumor grade and subtype classifications^1^, but also information on the tumor microenvironment (TME), such as the spatial organization of different types of cells. Cell spatial organization reveals cell growth patterns and the spatial interactions among different types of cells, which provide important insights into tumor progression and metastasis. A recent study by Cheng et al ^2^ developed an algorithm to segment the cell nucleus and extract the topological features for TME analysis in renal cell carcinoma (RCC), which improved the understanding of cell spatial organization and patient outcome in RCC. However, this study used an unsupervised approach that assigns an identified cell nucleus to one of the clusters without a clear definition, which hampered interpretation of the results. Recent studies^3, 4^ showed that the spatial organization and architecture of tumor-infiltrating lymphocytes (TIL) play important roles in the TME. However, these studies focused solely on recognizing lymphocytes and ignored other types of cells, which greatly limited exploration of the interactions among different types of cells. In this study, we developed a deep learning-based algorithm to examine standard H&E pathology images to automatically segment and classify different types of cell nuclei. This algorithm can be used as a tool to “computationally stain” different types of cell nuclei, in order to facilitate pathologists in examining tissue images and researchers in studying the TME from these standard clinical materials.

The major cell types in a malignant tissue include tumor cells, stromal cells, lymphocytes, and macrophages. Stromal cells are mainly connective tissue cells such as fibroblasts and pericytes. The interactions between tumor cells and stromal cells play an important role in cancer progression^5–7^ and metastasis inhibition^8^. Since the cell boundaries of tumor cells and stromal cells are often unclear in standard H&E stained lung cancer pathology images, we segmented and classified cell nuclei instead of whole cells. TIL are mainly white blood cells that have migrated into a tumor region. They are a mix of different types of cells, in which T cells are the most abundant population. The spatial organization of TIL has been associated with patient outcome and molecular profiles in multiple tumor types^9–12^. Macrophages are inflammatory cells, and inflammation in tumor niches has been reported as a prognostic marker and correlated with tumor progression^13–15^. Other tissues and cellular structures existing in the TME include blood vessels and necrosis. In this study, blood cells and karyorrhexis are segmented to represent blood vessels and necrosis, respectively, in order to quantify blood vessels and necrosis and study their interactions with tumor cells, stromal cells, lymphocytes and macrophages.

In this study, we developed a deep learning algorithm based on the Mask Regional Convolutional Neural Network (Mask-RCNN) architecture^16^. We trained this Mask-RCNN model using pathology images of lung adenocarcinoma (ADC) patients from the National Lung Screening Trial (NLST) study with the nuclei of tumor cells, stromal cells, lymphocytes, macrophages, blood cells and karyorrhexis manually labeled by expert pathologists. The model accuracy was validated in a different set of images. From the identified cell types and cell spatial locations, we derived cell spatial organization features to characterize the TME. In our analysis, we found that these TME-related image features were significantly associated with patient overall survival. Based on these image features, a prognostic model for lung ADC patients was developed from the NLST dataset. This model was independently validated in the pathology image data from The Cancer Genome Atlas (TCGA) lung ADC (LUAD) dataset, in which the predicted high-risk group showed significantly worse survival than the low-risk group (pv= 0.001), with a hazard ratio of 2.23 [1.37-3.65] after adjusting for clinical variables.

Furthermore, the image-derived TME features were significantly correlated with the gene expression of biological pathways. For example, transcription activation of both the T-cell receptor (TCR) and Programmed cell death protein 1 (PD1) pathways was positively correlated with the density of detected lymphocytes in tumor tissues, while expression of the extracellular matrix organization pathway was positively correlated with the density of stromal cells.

In this study, we developed a Mask-RCNN algorithm for nuclei segmentation and cell classification as a tool to study the tumor morphological microenvironment using tissue pathology images in lung ADC. In order to facilitate usage of this deep learning tool, a user-friendly web portal has been developed and can be accessed at http://lce.biohpc.swmed.edu/maskrcnn/analysis.php. Although this tool was developed in lung ADC pathology images, our results showed that the Mask-RCNN method can be adapted and applied in head and neck cancer, breast cancer and lung cancer squamous cell carcinoma pathology image datasets. The web portal also provides functions to facilitate researchers in adapting the Mask-RCNN model for other types of cancers.

## 2 METHODS

### 2.1 Dataset

Pathology images that support the findings of this study are available online in NLST (https://biometry.nci.nih.gov/cdas/nlst/) and The Cancer Genome Atlas Lung Adenocarcinoma (TCGA-LUAD, https://wiki.cancerimagingarchive.net/display/Public/TCGA-LUAD). mRNA expression data for the TCGA dataset are available online at http://firebrowse.org. The H&E-stained pathology images together with the corresponding clinical data were obtained from the NLST and TCGA Lung ADC cohorts: 208 40X pathology images for 135 lung ADC patients were acquired from the NLST dataset, and 431 40X pathology images for 372 lung ADC patients were acquired from the TCGA LUAD dataset (there could be multiple pathology images for a single patient). A specialized lung cancer pathologist, Dr. Lin Yang, labeled the Region of Interest (ROI) for each of the pathology images (Figure 1). Another lung cancer pathologist, Dr. Adi Gazdar, confirmed the labelling. Clinical characteristics of the patients in this study are summarized in Supplemental Table 1.

**Figure 1.**
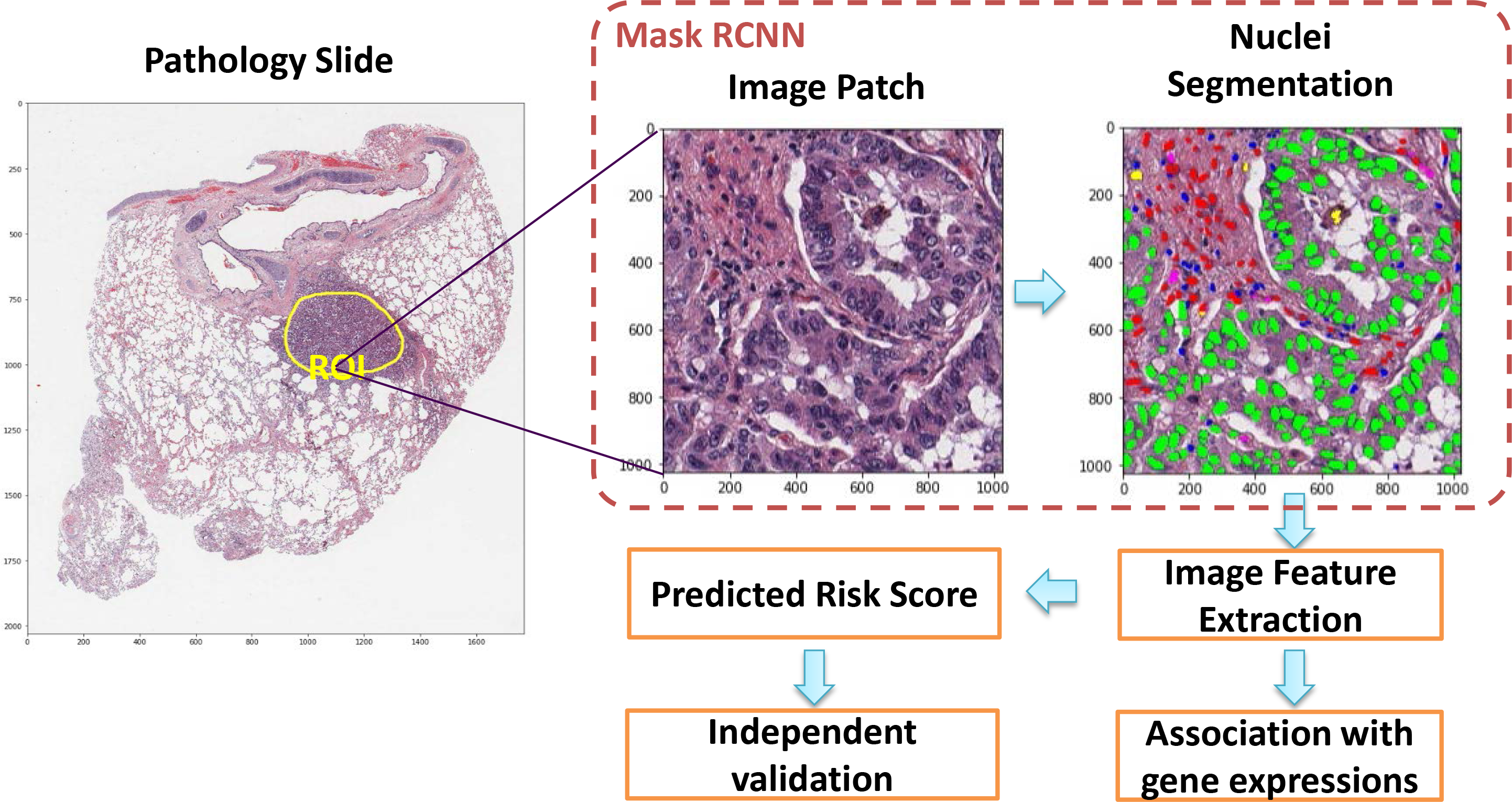
Flow chart of pathology image analysis pipeline in this study. Mask RCNN: Mask Regional-Convolutional Neural Network.

### 2.2 Nuclei segmentation using Mask-RCNN

#### 2.2.1 Training, validation, and testing sets preparation

In order to construct the training set for the Mask-RCNN algorithm, 127 image patches (500 × 500 pixels) from 39 pathological ROIs (Figure 1) were extracted from the NLST dataset. In these patches, different types of cell nuclei were labeled. All the pixels with tumor nuclei, stromal nuclei, lymphocyte nuclei, macrophage nuclei, red blood cells, and karyorrhexis were labeled according to their categories and all the remaining pixels were considered “other”. These labels, also collectively called the “mask”, were then used as the ground truth to train the Mask-RCNN model. The labeled images were randomly divided into training, validation, and testing sets. To ensure independence among these datasets, image patches from the same ROI were assigned together. More than 12,000 cell nuclei were included in the training set, while 1227 and 1086 nuclei were included in the validation and testing sets, respectively.

#### 2.2.2 Training process

A deep learning model was developed using the Mask-RCNN architecture. The pre-trained model was fine-tuned on our training dataset from the NLST study. Images were standardized (centered and scaled to have zero mean and unit variance) for each RGB channel. To increase generalizability and avoid bias from different H&E staining conditions, we performed extensive augmentations on the image patches. In particular, random projective transformations were applied to images and their corresponding masks; each image channel was randomly shifted using linear transformation. For the training process, the batch size was set to 2, the learning rate was set to 0.01 and decreased to 0.001 after 500 epochs, the momentum was set to 0.9, and the maximum number of epochs to train was set to 1000. In the validation set, the model trained at the 707^th^ epoch reached the lowest loss. This model was selected and used in the following analysis to avoid overfitting. Python (version 3.5.2) and python libraries (Keras, version 2.1.5; openslide-python, version 1.1.1; tensorflow-gpu, version 1.8.0) were used^17^.

#### 2.2.3 Segmentation performance evaluation

Since the Mask-RCNN model simultaneously segments and classifies cell nuclei, three criteria were used to evaluate the segmentation performance in the validation and testing datasets, respectively. First, detection coverage was calculated as the ratio between the detected nuclei and the total ground truth nuclei. Each ground truth nucleus was matched to a segmented nucleus, which generated the maximum Intersection over Union (IoU). If the IoU for a ground truth nuclei was > 0.5, this nuclei was labeled “matched”; otherwise it was labeled “unmatched”. Second, nuclei classification accuracy was determined for the matched nuclei by comparing the predicted nucleus type with the ground truth. Third, segmentation accuracy was evaluated by the IoUs, which were calculated for each detected nucleus and averaged in different nuclei categories.

### 2.3 Image feature extraction to describe nuclei composition and organization

In order to make the Mask-RCNN model more computationally efficient while retaining a good representation of each ROI, instead of applying the model to the whole slide, 100 image patches (1024 × 1024 pixels) were randomly sampled and analyzed for each pathologist-labeled ROI (Supplemental Figure 1). These 100 image patches provided good coverage of each ROI. Nuclei were then segmented and classified through the Mask-RCNN model developed from this study (Figure 1). In order to characterize the spatial organization of cells using a graph, we calculated the centroids of nuclei and used them as vertices to construct a Delaunay triangle graph for each image patch. The Delaunay triangle graph connects nuclei into a graph, and the number of connections and the average length (i.e. spatial distance) between two types of nuclei summarize the spatial organization of different types of cell. Since 6 nucleus categories were included in this study, the edges of the graph were classified into 21 categories [i.e. 6 × (6−1)/2 = 21] according to their vertex pairs. For each image patch, the number of connections (i.e. edges) for different categories was counted (which added up to 21 features), the lengths of the connections were averaged for each edge category (yielding another 21 image features), and the density of each type of nucleus was calculated (yielding 6 image features). In total, 48 image features were extracted. The image features were averaged across the 100 patches for each ROI in the pathology image. When 2 or more pathology slides were available for 1 patient, the features from the slides were averaged for each patient. Thus, in total 48 image features were extracted for each patient, in both the NLST and TCGA datasets.

### 2.4 Prognostic model development and validation

Overall survival, defined as the date of diagnosis till death or last contact, was used as the response variable for survival analyses. A prognostic model for overall survival was developed in the lung ADC patients in the NLST dataset and independently validated on the TCGA LUAD dataset. Given a set of image-derived TME features for each patient, the prognostic model will assign a risk score for the patient, with a higher risk score indicating worse prognosis. Based on the risk scores, the patients in the TCGA LUAD cohort were dichotomized into predicted high- and low-risk groups using the median risk score as a cutoff. The survival curves of the predicted high- and low-risk groups were estimated using the Kaplan-Meier method. The survival differences between predicted high- and low-risk groups were compared using a log-rank test. A multivariate Cox proportional hazard model was used to determine the prognostic value of predicted risk groups (using image-derived TME features) after adjusting for other clinical characteristics, including age, gender, smoking status, and stage. R software, version 3.4.2, and R packages (survival, version 2.41-3; glmnet, version 2.0-13; spatstat, version 1.55-1) were used^18, 19^. The results were considered significant if the two-tailed p value <0.05.

### 2.5 Association analysis between image features and gene expression of biological pathways

Gene expression data of 372 patients from the TCGA LUAD dataset were downloaded and preprocessed. The correlation between mRNA expression levels and image-derived TME features was evaluated using Spearman rank correlation. Gene set enrichment analysis (GSEA) was performed for each TME feature. All gene sets from the Reactome database were used^20^. For multiple testing correction, Benjamini-Hochberg (BH)-adjusted p values were used to detect significantly enriched gene sets. Gene sets with BH-adjusted two-tailed p values < 0.05 were regarded as significantly enriched. R packages Hmisc (version 4.1-1), fgsea (version 1.4.1), and gplots (version 3.0.1) were used^21^.

## 3 RESULTS

### 3.1 Mask-RCNN simultaneously and accurately classifies and segments cell nuclei

The developed Mask-RCNN model segments and classifies individual nuclei at the same time (Supplemental Figure 2). Figure 2 demonstrates some of the segmentation and classification results. In total, the segmented cell nuclei were classified into six categories: tumor cell, stromal cell, lymphocyte, macrophage, red blood cell, and karyorrhexis, and all the remaining structures or spaces were considered background. Different nuclei were colored according to the predicted categories (Figure 2). For detected objects, the overall classification accuracy was 85% and 85% in the validation set and the testing set, respectively, while the accuracy for tumor nuclei was 88% in validation and 90% in testing, respectively (Supplemental Figure 3). It is noteworthy that the developed Mask-RCNN model can be applied to the entire digital pathology image to generate a cell spatial organization map across the whole slide, where tumor region and lymphocyte infiltration areas are clearly illustrated (Figure 3).

**Figure 2.**
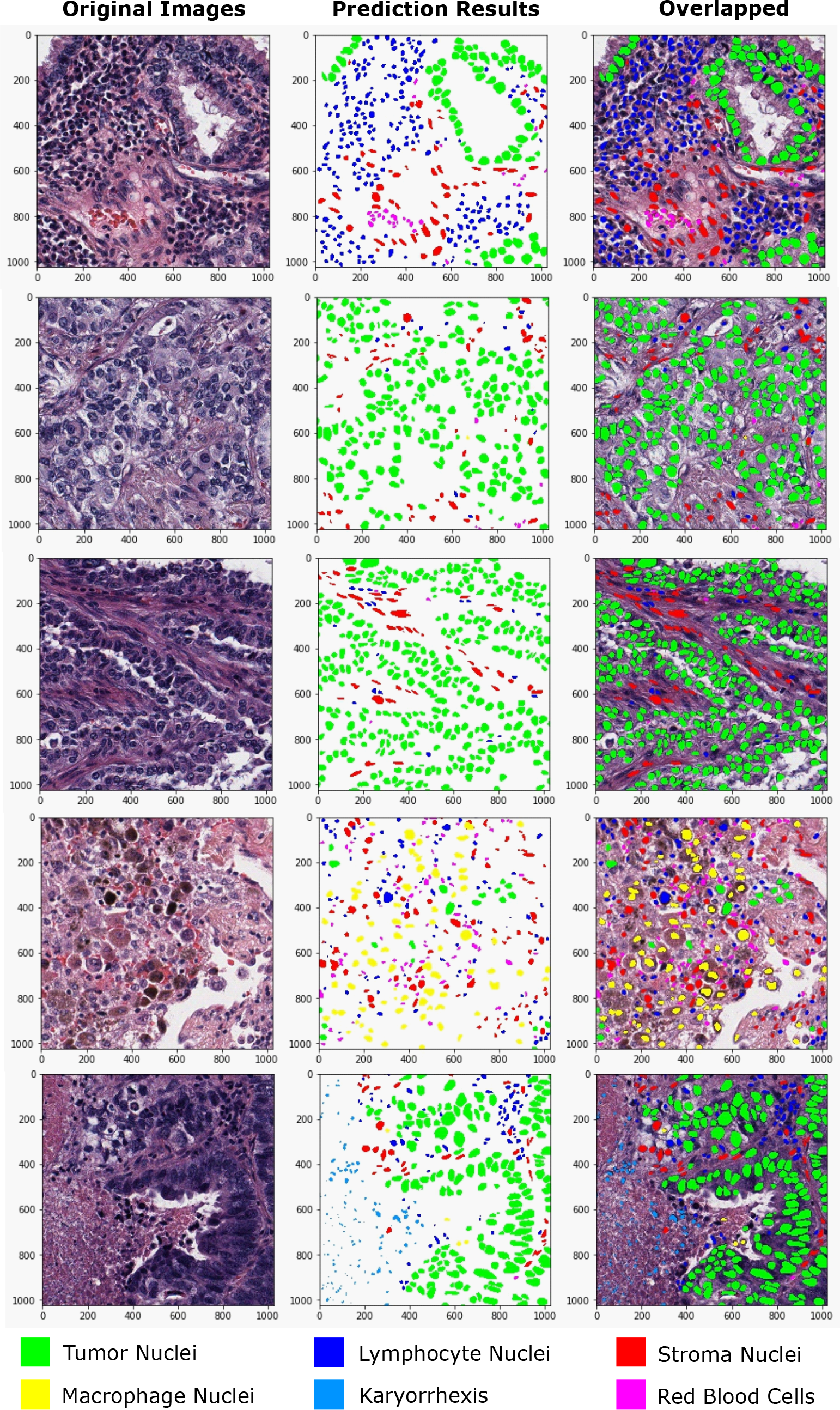
Mask-RCNN-based nuclei segmentation and classification results in lung ADC pathological images.

**Figure 3.**
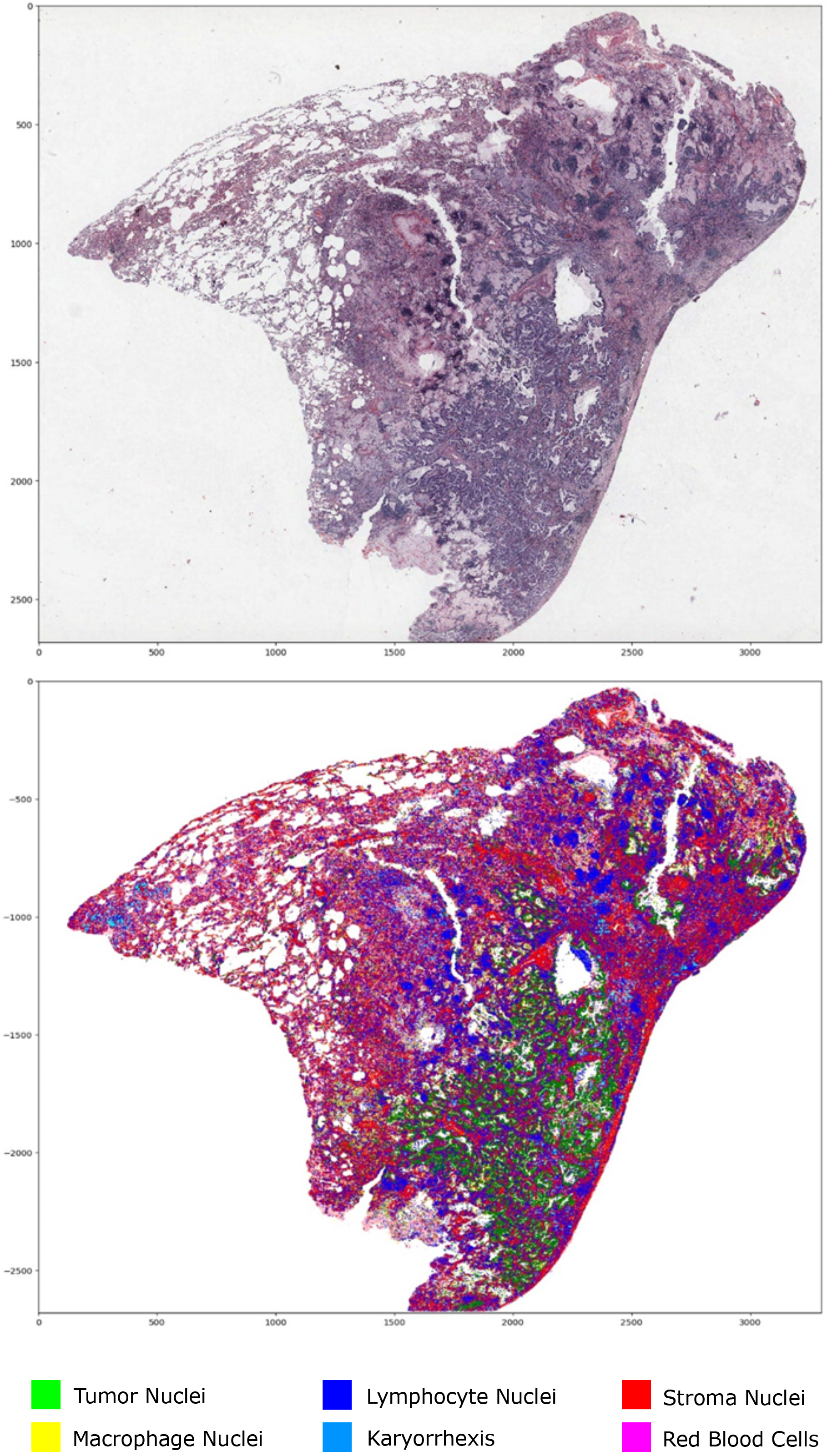
Nuclei segmentation and classification across whole slide image. Upper panel: Original pathology image. Lower panel: detected and classified nuclei overlay on top of the original pathology image.

### 3.2 Prognostic value of nuclei composition and organization in the TME

A Delaunay triangle graph was constructed for each image patch^2^ to extract topological features from nuclei spatial organization to characterize the TME (Supplemental Figure 4). The nuclei and edges were counted and edge lengths averaged to yield 48 image features (see **Methods Section**). Supplemental Table 2 summarizes the TME features that were significantly correlated with survival outcome in univariate analysis. It shows that higher karyorrhexis density, more karyorrhexis-karyorrhexis connections and more karyorrhexis-red blood cell connections were associated with worse survival outcome, which was expected as these features indicate a higher rate of tumor necrosis. Furthermore, higher stromal nuclei density and more stromal-stromal connections were associated with better survival outcome, which agreed with our current knowledge that more stromal tissues corresponds to better prognosis.

A prognostic model based on the image features was developed in the NLST dataset and then independently validated in the TCGA LUAD dataset. Figure 4A shows the survival curves of the predicted high and low-risk groups in the TCGA cohort, where the patients in the predicted high-risk group show significantly worse survival than those in the predicted low-risk group (log-rank test, p value = 0.0011). Furthermore, the risk group defined by the TME features serves as an independent prognostic factor (high- vs. low-risk, Hazard Ratio = 2.23, 95% Confidence Interval = 1.37-3.65, and p value=0.0013), after adjusting for clinical variables, including age, gender, smoking status, and stage (Figure 4B).

**Figure 4.**
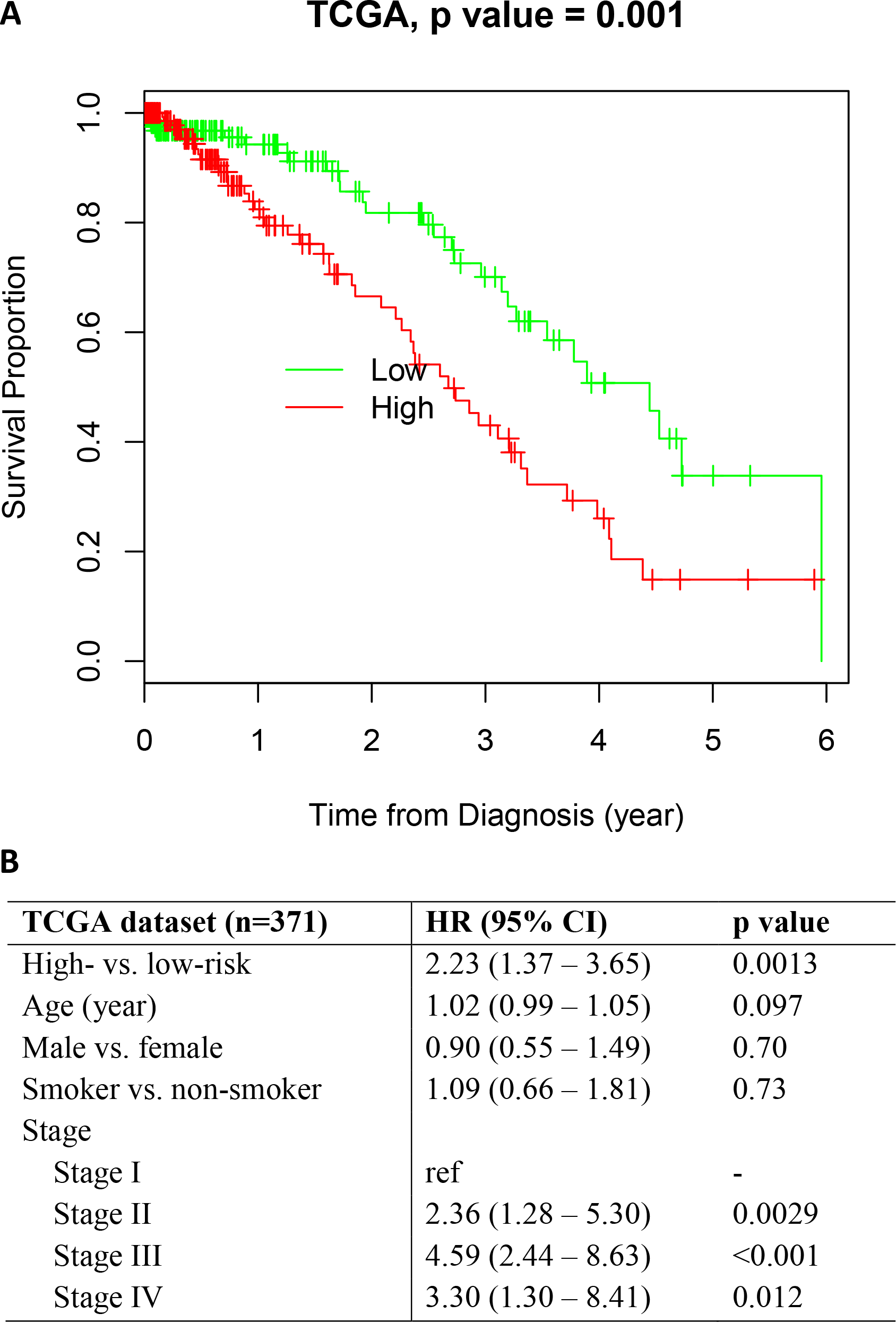
Prognostic value of the TME feature-based prognostic model. **(A)** K-M plot of predicted high- and low-risk groups in the TCGA dataset. Log-rank test, p-value = 0.001. **(B)** Multivariate survival analysis of predicted risk group adjusted by potential confounders. CI, confidence interval; HR, hazard ratio.

### 3.3 Association between image features and transcriptional activity of biological pathways

GSEA was performed to identify the biological pathways whose mRNA expression profiles were significantly correlated with image-derived TME features in the TCGA dataset. Figure 5A-D shows examples of these biological pathways. For example, the transcription activation of both the T-cell receptor (TCR) and Programmed cell death protein 1 (PD1) pathways was positively correlated with lymphocyte density in the tumor tissue (Figure 5A). This observation is consistent with previous reports that genes involved in the TCR and PD1 pathways are expressed in immune cells^22, 23^. In addition, expression of the extracellular matrix organization gene set, for which fibroblasts act as an important source^24^, was positively correlated with stromal cell density in tumor tissue (Figure 5B). In a negative control experiment where we randomly shuffled the patient IDs and repeated the same analysis, such correlation was no longer observed (Supplemental Figure 5).

**Figure 5.**
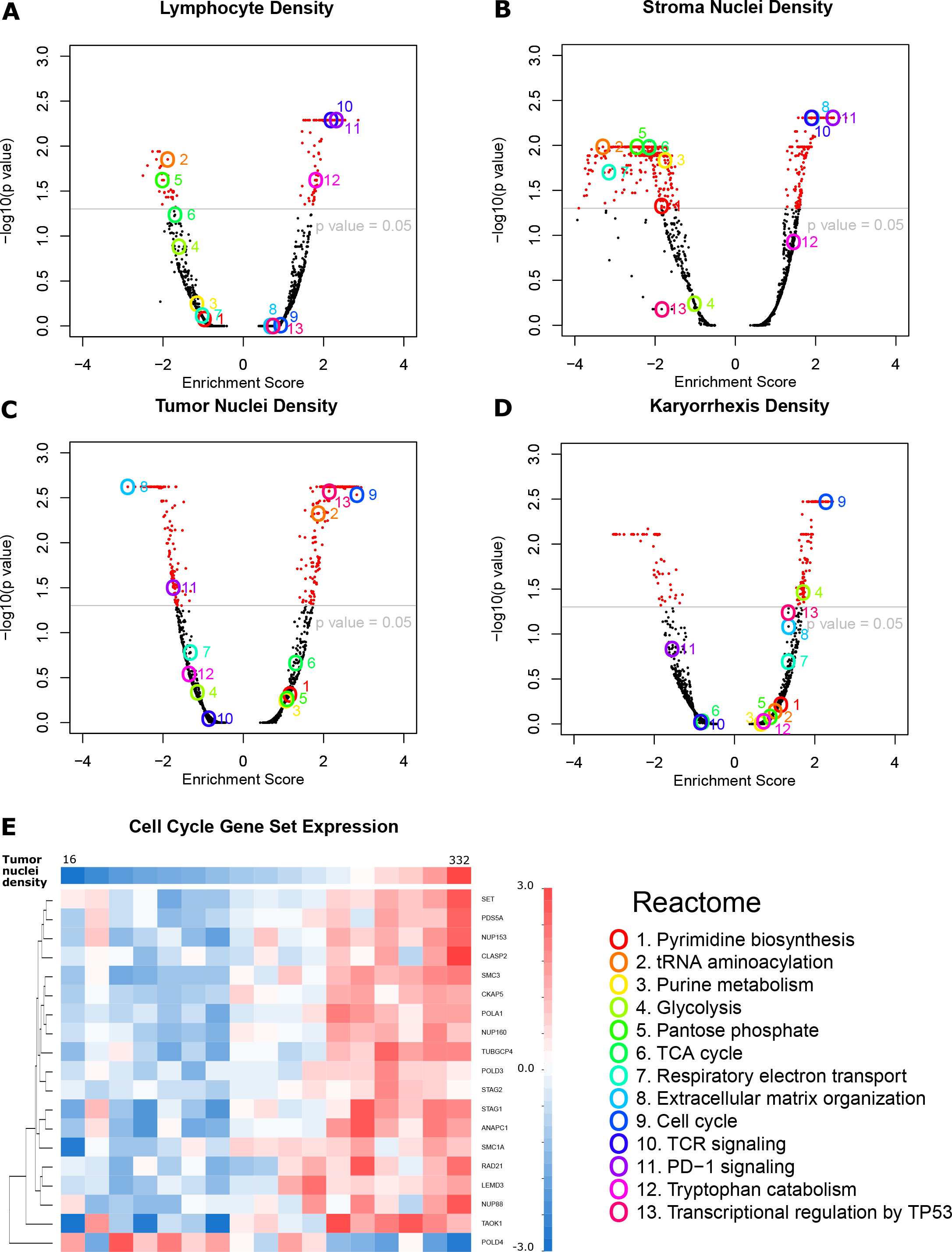
Correlation between image features and mRNA expression in tumor. **(A-D)** Volcano plots of gene set enrichment analysis results correlating mRNA expression level with tumor nuclei density (A), stromal nuclei density (B), lymphocyte density (C), and karyorrhexis density (D) respectively. Thirteen interesting gene sets are highlighted. **(E)** To look into the significantly correlated gene sets, an example heatmap shows that most mRNA expression levels in the cell cycle gene set are positively correlated with tumor nuclei density in tumor issue. Only genes with p value < 0.001 in Spearman rank correlation with tumor cell number are shown. Patients are grouped according to tumor cell density showing on the top row.

Furthermore, GSEA analysis showed that the cell cycle pathway was significantly enriched with genes whose expression levels were correlated with both the tumor nuclei density (Figure 5C) and karyorrhexis density in tumor issue (Figure 5D). To look into the relationship between tumor cell density and the gene expression of the cell cycle pathway, we grouped and sorted the patients in the TCGA LUAD cohort according to their tumor nuclei density. Figure 5E shows, for each patient group in the TCGA LUAD dataset, the average expression levels of genes within the cell cycle pathway and whose expression levels are significantly (p value <0.001) correlated with tumor nuclei density. Positive correlations between gene expression and tumor nuclei density can be observed for most of the cell cycle-related genes, except for one gene, POLD4, which showed an inverse trend. Most of the genes in the cell cycle pathway have higher expression in tumors with higher tumor nuclei density (may be a higher grade of tumor), while POLD4 shows the opposite pattern. This pattern of POLD4 compared with other genes in the cell cycle gene set is consistent with a previous study of lung cancer^25^: while most cell cycle genes were upregulated in lung cancer, POLD4 is usually downregulated.

### 3.4 Webserver for publically accessible pathological image segmentation model

In order to facilitate usage of the Mask-RCNN model developed in this study, we also developed an online tool (http://lce.biohpc.swmed.edu/maskrcnn/analysis.php) for this deep learning-based nuclei segmentation and classification model (Figure 6). This tool requires only a pathology image (or a patch from the image) as the input (Figure 6A). Each uploaded input image will be assigned a job ID (Figure 6B). The segmentation results will be automatically displayed and the spatial coordinates of each nucleus can be downloaded as an Excel table (Figure 6C). In order to assist researchers in using this tool to study TME-related features for other cancer types, we also provide a function to automatically generate a mask for other cancer types. The newly generated segmentation mask can greatly reduce the manual work of creating the training sets for other cancer types, and thus accelerate the development of applications for pathology image analysis. Results show that the Mask-RCNN method can be adapted and applied in head and neck cancer, breast cancer and lung cancer squamous cell carcinoma pathology image datasets in TCGA (Supplemental Figure S6, S7 **and** S8).

**Figure 6.**
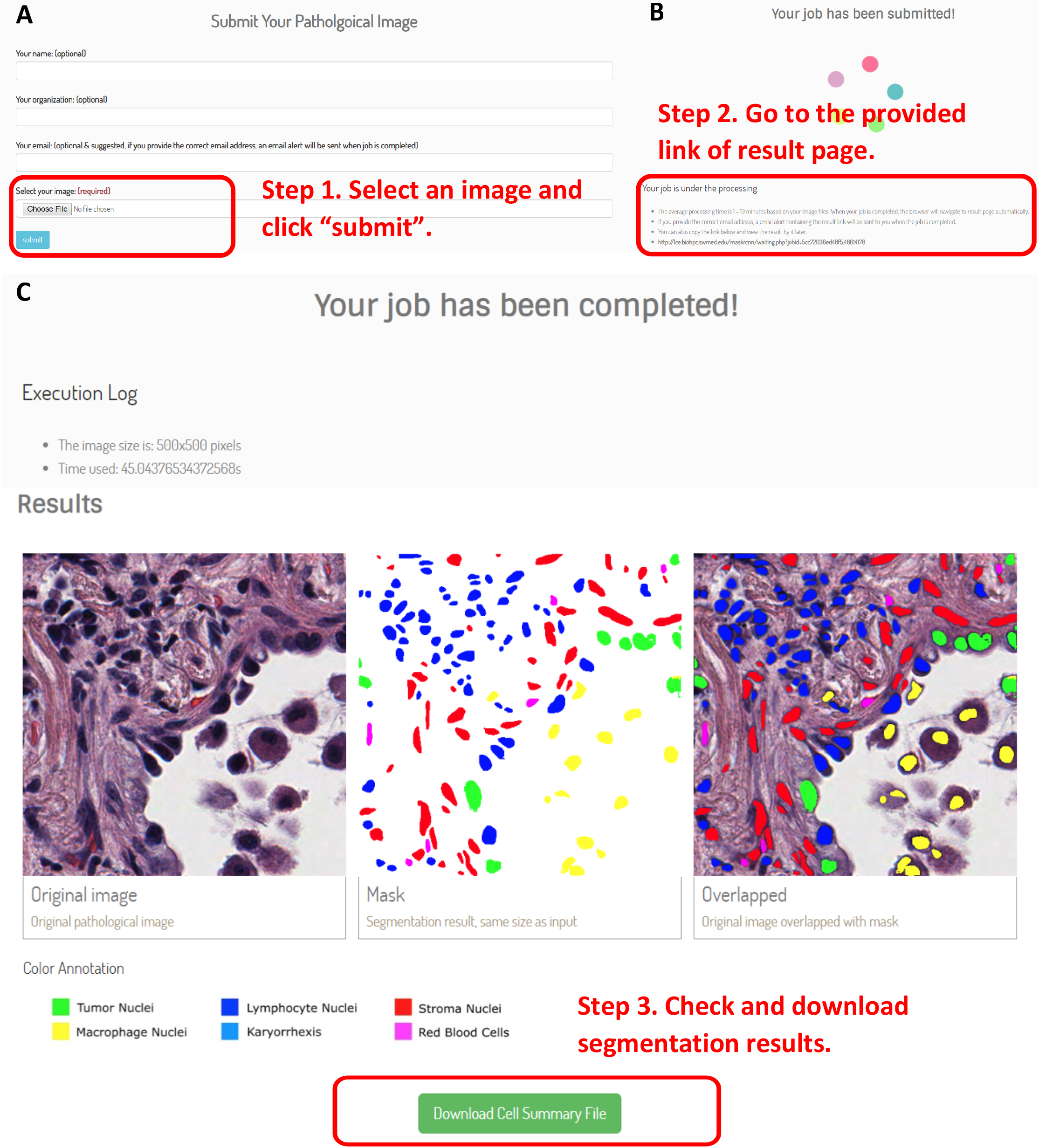
An online tool for nuclei segmentation and classification in pathology image. Nuclei segmentation and classification results can be automatically generated from pathology images. The spatial locations and cell types of individual cells can be downloaded as an Excel table.

## 4 DISCUSSION

In this study, we developed a deep learning-based analysis tool to study the TME using standard H&E stained pathology images. This tool successfully visualized and quantified the spatial organization of tumor cells, stromal cells, lymphocytes, inflammatory cells, red blood cells and karyorrhexis in the tumor tissues of lung ADC patients. The topological features of cell spatial organization were used to characterize the TME. Our results showed that these features were associated with patient survival outcome and the gene expression of biological pathways. From these image-derived TME features, we developed a prognostic model for lung ADC patients and independently validated it in another lung ADC patient cohort. The prognostic model predicts patient survival independent of other clinical variables in the validation cohort.

Several previous studies have tried to analyze the TME and discovered prognostic image features. However, these studies involved time-consuming hand-labeling by pathologists^26–28^. In contrast, we developed a fully automated and objective nuclei segmentation and classification strategy. In addition, this deep learning-based method enables segmentation of nuclei within a whole slide image. Since the number of cells in a whole slide image could be tremendous (~2,000,000 on average), manually labelling all of them is impractical. Thus, this deep-learning method empowers quantification of the TME across the whole slide image. Furthermore, although developed in lung ADC, this method can be easily generalized to other cancer types by retraining the model using the tools provided by our web portal.

In pathology image analysis, three-dimensional tissue structures are captured as two-dimensional images, and the cell nuclei may “touch” and “overlap” with each other in the resulting images. This is one of the major challenges for nuclei segmentation in pathology image analysis. In this study, we developed a Mask-RCNN-based deep learning model to segment and classify different types of cell nuclei in order to study the spatial interactions among different cell types and tissue structures. Compared with other image segmentation algorithms, the Mask-RCNN method has several advantages: First, it segments and classifies nuclei at the same time, while traditional nuclei segmentation algorithms relying on color deconvolution cannot classify cell types^2, 29^. Second, by using extensive color augmentation during the training process, it adapts to different staining conditions, which makes the algorithm more robust and allows us to avoid the time-consuming color normalization steps ^30^. Third, compared with other popular semantic image segmentation neural networks such as Fully Convolutional Network (FCN) and Deeplab^31, 32^ that classify each pixel, Mask-RCNN is intrinsically an instance segmentation algorithm that detects an object bounding box first and assigns pixels as foreground or background within this bounding box^16^. In summary, Mask-RCNN provides a new solution to segmenting closely clustered nuclei in tissue pathology images.

The associations between the extracted TME features and patient prognosis were evaluated in this study. Karyorrhexis, a representative of necrosis, has been reported as an aggressive tumor phenotype in lung cancer^33^. Consistently, the density of karyorrhectic cells and numbers of karyorrhexis-karyorrhexis edges were shown as negative prognostic factors in this study. On the other hand, the density of stromal cells and the numbers of stromal cell-stromal cell edges were positive prognostic factors, which is consistent with a recent report on lung ADC patients^8^. These consistencies indicate the validity of this MaskRCNN-based deep neural network and the potentiality of using cell organization features as novel biomarkers for clinical outcomes.

Gene expression patterns have been widely used to study the underlying biological mechanisms of different tumor types and subtypes^35, 36^; moreover, genes with abnormal expression could become potential therapeutic targets of cancers ^37, 38^. However, traditional transcriptome profiling is usually done in bulk tumor^35, 39^, which contains multiple cell types, such as stromal cells and lymphocytes, in addition to tumor cells. This bulk tumor-based sequencing could blur or diminish the mRNA expression changes arising from a single cell type or from different cell compositions in the TME. Currently, the relationship between the transcription activities of biological pathways and the TME remains unclear. In this study, the image-derived TME features show interesting correlations with the transcriptional activities of biological pathways. For example, gene expression levels of TCR and PD-1 pathways were positively correlated with the density of lymphocytes detected from tumor tissues. This indicates the image-derived TME features may be used to study or predict immunotherapy response, since several promising cancer immunotherapies rely on activation of tumor-infiltrated immune cells and blocking immune checkpoint pathways^23, 34^. In addition, the gene expression extracellular matrix organization pathway is associated with the density of stromal cells in tumor tissues. Since traditional transcriptome sequencing is done in bulk tumor, accurate cell composition derived from pathology images could help to improve the evaluation of gene expression for each individual cell type. Moreover, the correlation between image features and transcriptional patterns of biological pathways hints at the potential usage of image features to study tumor bioprocesses, including cell cycle and metabolism status.

There are some limitations to this Mask-RCNN model. First, information on the individual nuclei, such as nucleus shape and size, was not considered since this study focused on nuclei organization. Morphological and intensity features of nuclei have been reported as prognostic factors, which can be automatically extracted using this nuclei segmentation algorithm^40^. Second, some special structures, such as bronchi and cartilage, were not included in this algorithm. This study handled this problem by avoiding such structures during ROI annotation. However, a more comprehensive training set would be desirable for whole slide analysis.

## Code availability

The code is available upon request. We will share the code through Github once this manuscript has been accepted.

## Author contributions

S.W. and G.X. designed the study. S.W. and R.R. implemented and trained the neural network model. S.W., L.C. performed genomic analysis. S.W., R.R., D.M.Y., L.C., L.X., T.W., Y.X., A.G., J.M. and G.X. analyzed the data and wrote the manuscript. L.Y. labeled the data. All authors commented on the manuscript.

## Acknowledgement

We thank Jessie Norris for helping us to edit this manuscript.

## Supplemental Material

**Supplemental Table 1.**
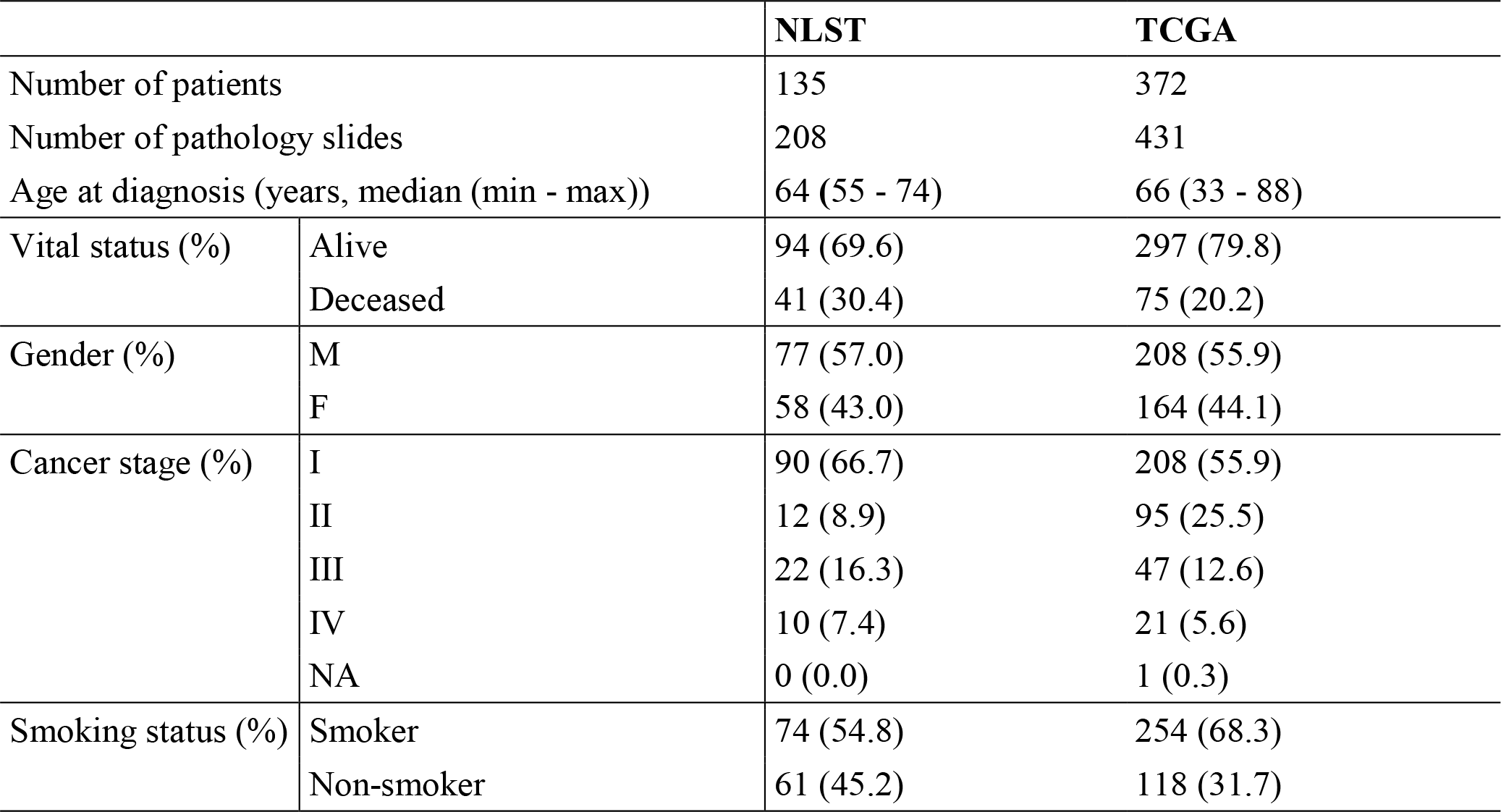
Patient characteristics for the National Lung Screening Trial (NLST) and The Cancer Genome Atlas (TCGA) datasets.

**Supplemental Table 2.**
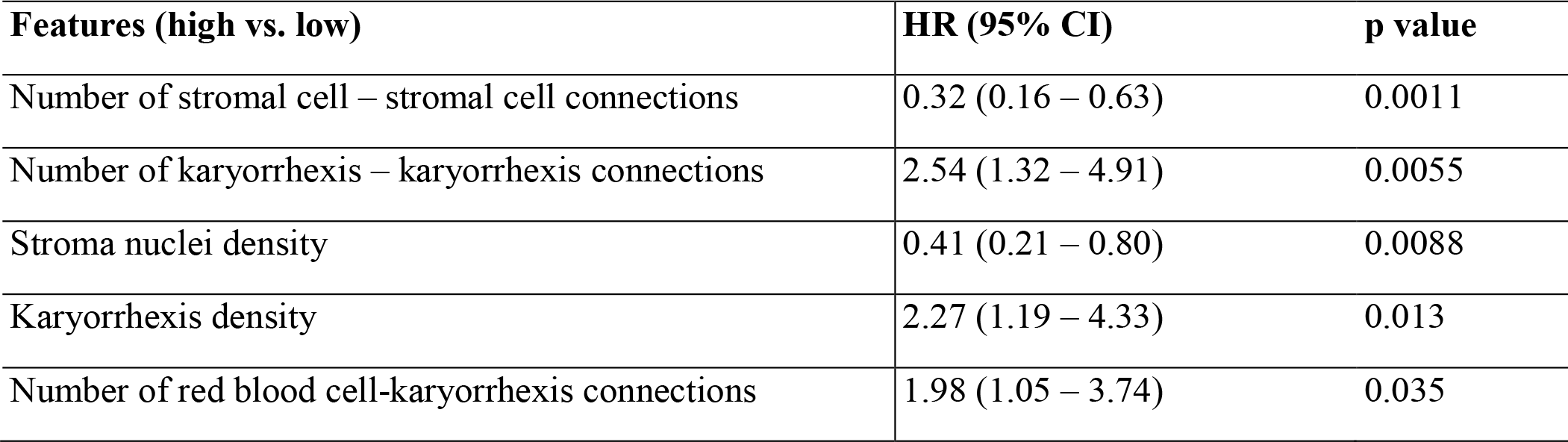
Image features that are significantly associated with patient overall survival outcome in univariate survival analysis in the National Lung Screening Trial (NLST) dataset. Features are dichotomized by the median value. P values are calculated using Ward test. CI, confidence interval; HR, hazard ratio.

**Supplemental Figure 1.**
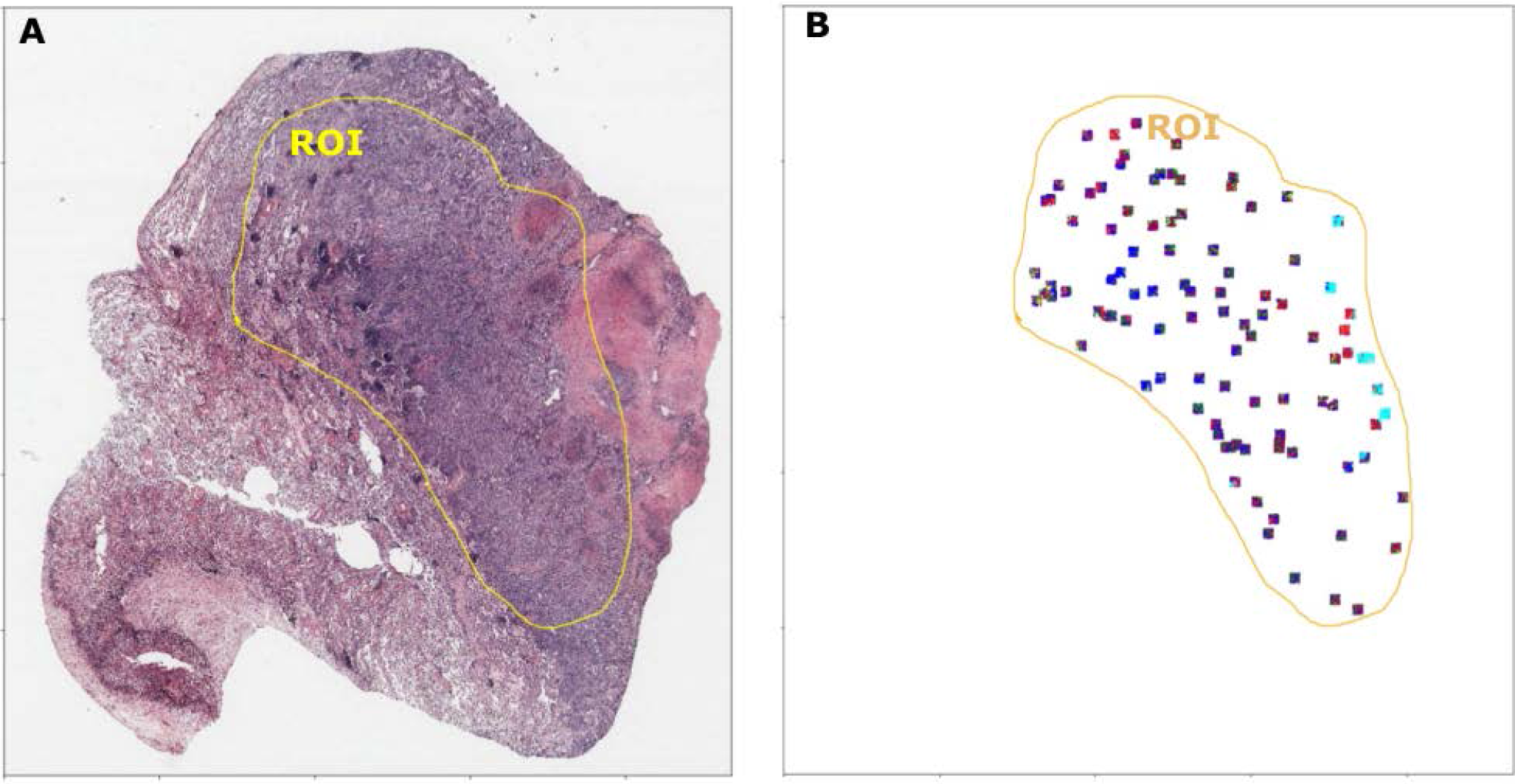
Illustration of 100 patches randomly sampled from the Region of Interest (ROI). **(A)** A pathology image with ROI labelled by pathologists (yellow line). **(B)** 100 patches are randomly sampled within the ROI. Each square represents a 1024 × 1024 pixel image patch.

**Supplemental Figure 2.**
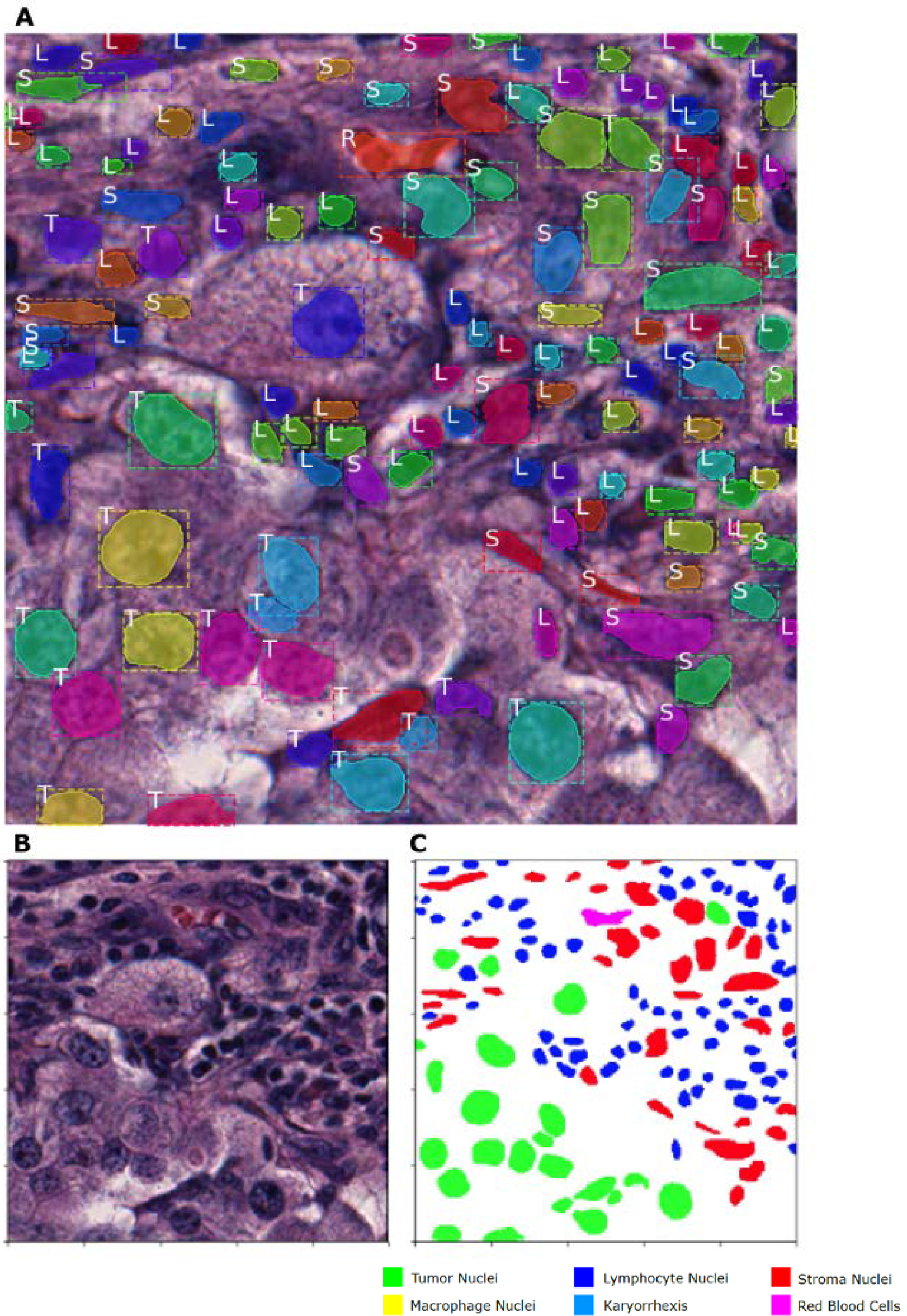
Illustration of instance segmentation using the developed Mask-RCNN model. **(A)** Each nucleus has a bounding box (dashed lines). Nucleus segmentation is made within this bounding box. The class of the nucleus is also predicted at the same time as the segmentation, and the nuclei class is labeled on the top left of its bound box. (T: tumor nuclei; S: stromal nuclei; L: lymphocyte nuclei; R: red blood cell; M: macrophage nuclei and K: karyorrhexis) **(B)** Original image patch (500 × 500 pixels). **(C)** For better viewing, the nuclei are colored according to their categories.

**Supplemental Figure 3.**
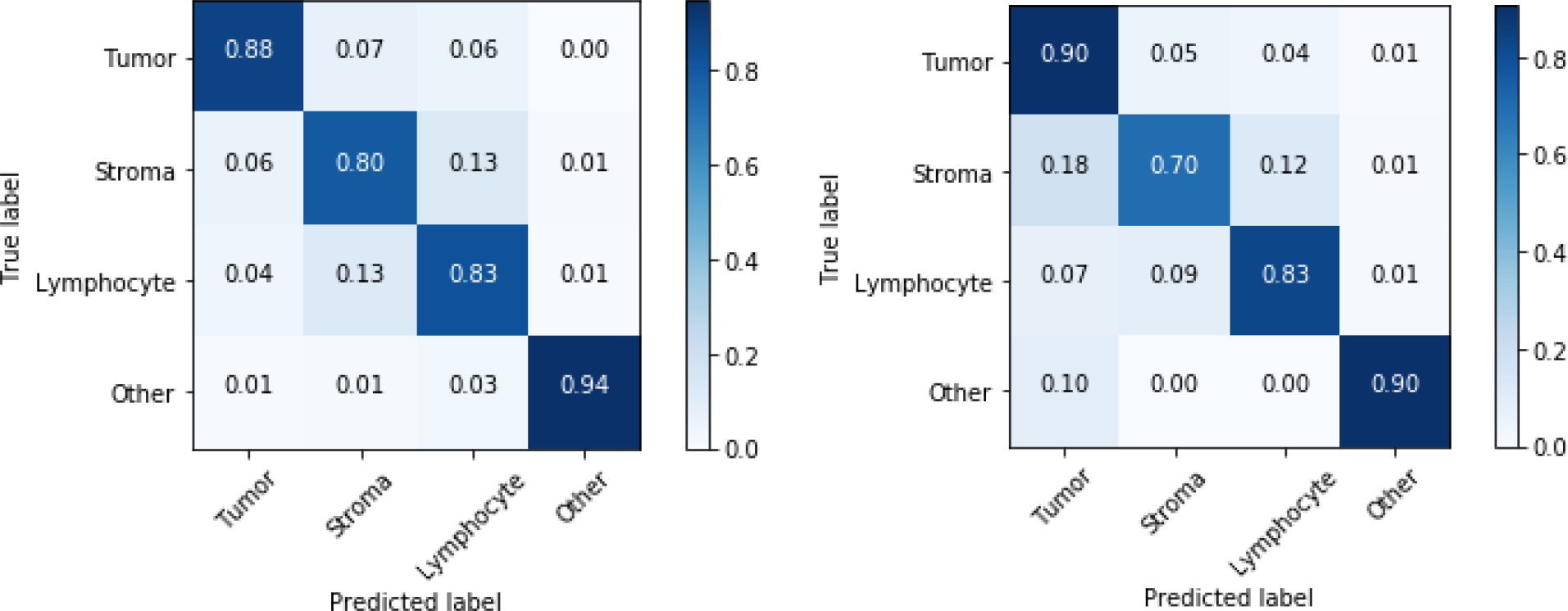
Confusion matrix between true labels and predicted labels, in the validation set (left) and testing set (right) respectively.

**Supplemental Figure 4.**
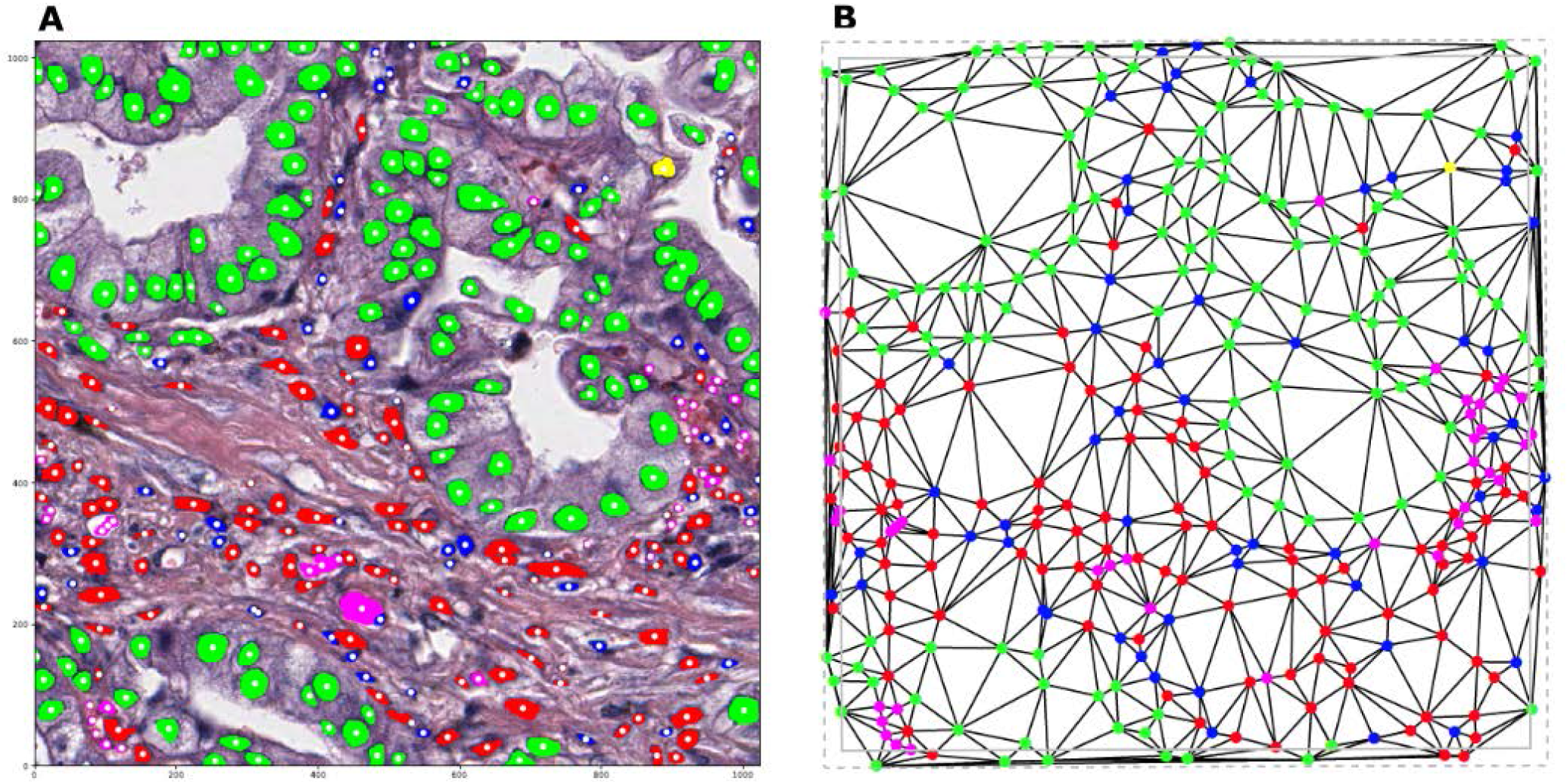
Illustration of Delaunay triangle construction. **(A)** Nuclei segmentation results using deep learning. Centroids of detected nuclei are marked in white. **(B)** A Delaunay triangle graph is constructed using nuclei centroids as vertices. To remove edge effect, when extracting graph properties, only edges with both ends within the solid gray square (976 × 976 pixels) are counted. The dotted gray square (1024 × 1024 pixels) plots the boundary of the input image patch. Green, tumor nuclei; red, stroma nuclei; blue, lymphocyte nuclei; yellow, macrophage.

**Supplemental Figure 5.**
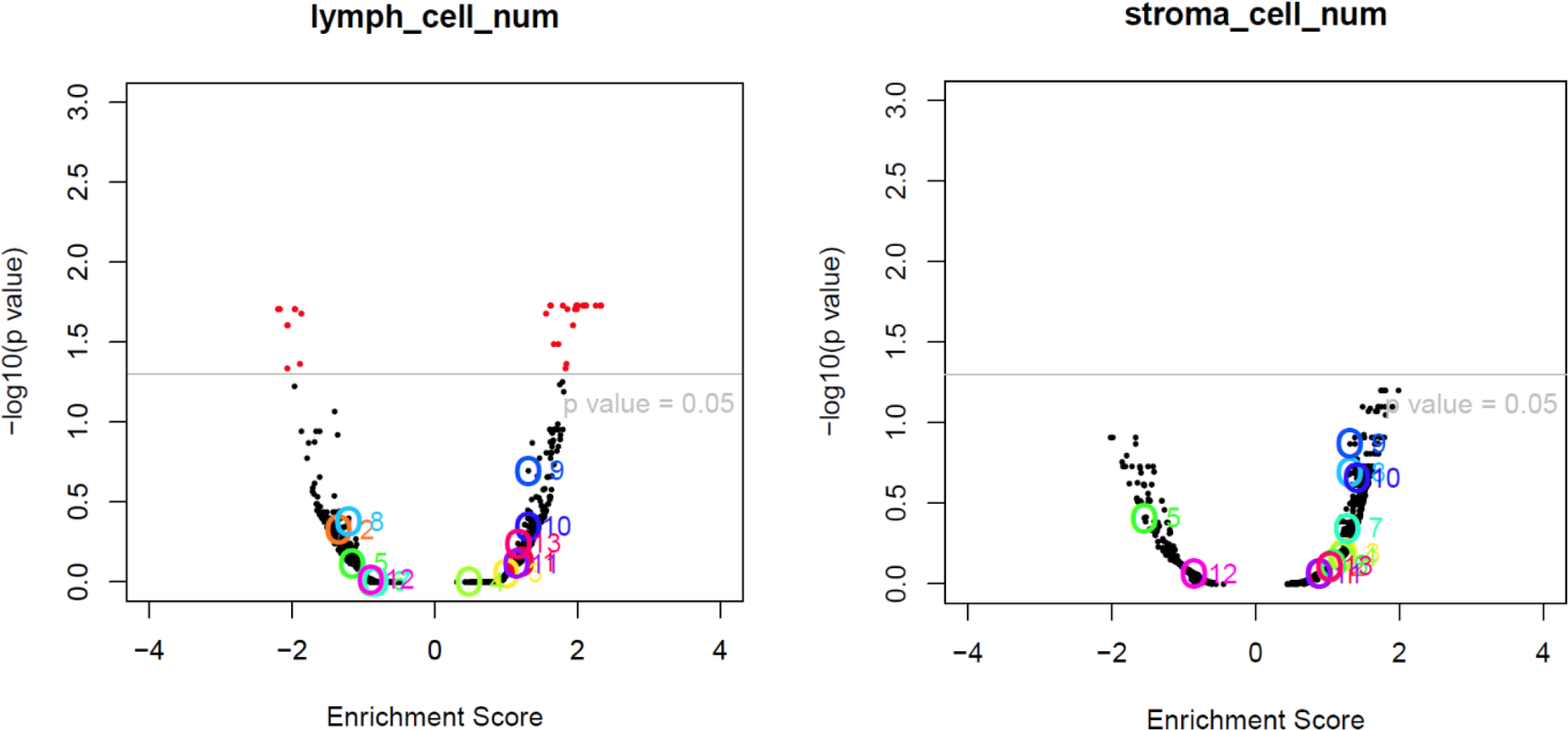
Example of gene set enrichment analysis (GSEA) results correlating mRNA expression with image-derived stromal cell density and lymphocyte density while patient IDs are randomly shuffled as a negative control experiment.

**Supplemental Figure 6.**
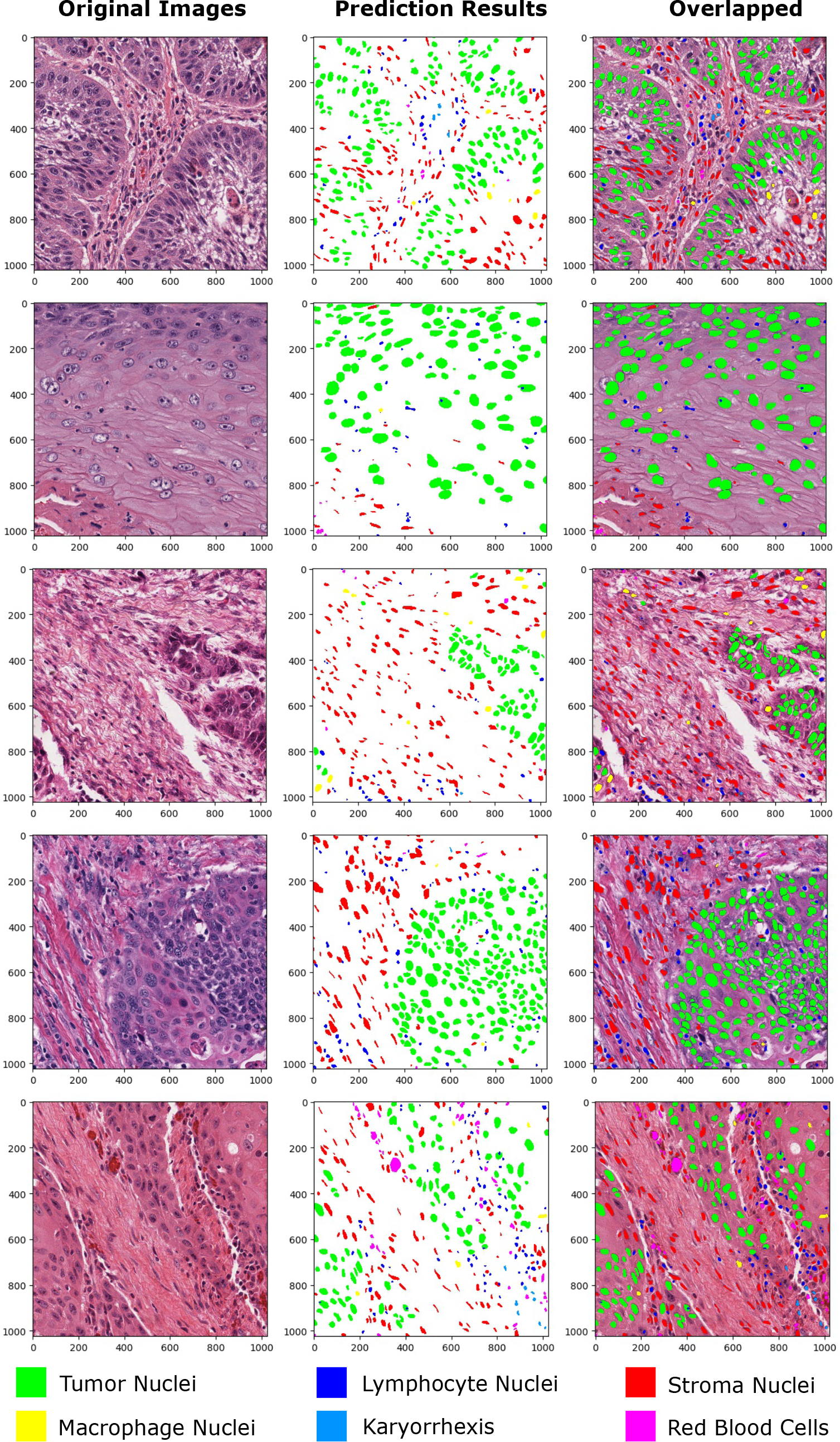
Deep learning based nuclei segmentation results in TCGA head and neck cancer pathology images.

**Supplemental Figure 7.**
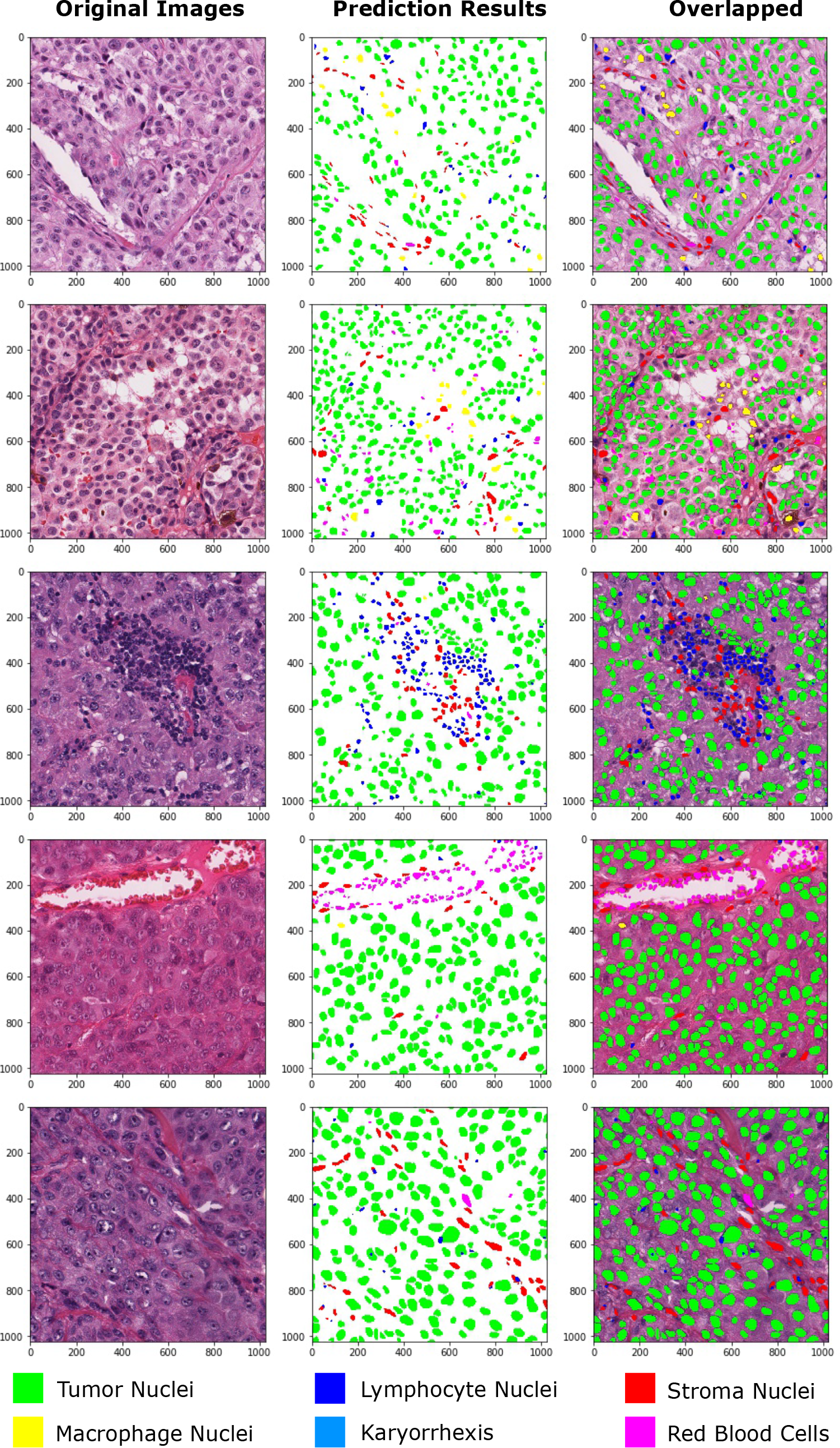
Deep learning based nuclei segmentation results in TCGA breast cancer pathology images.

**Supplemental Figure 8.**
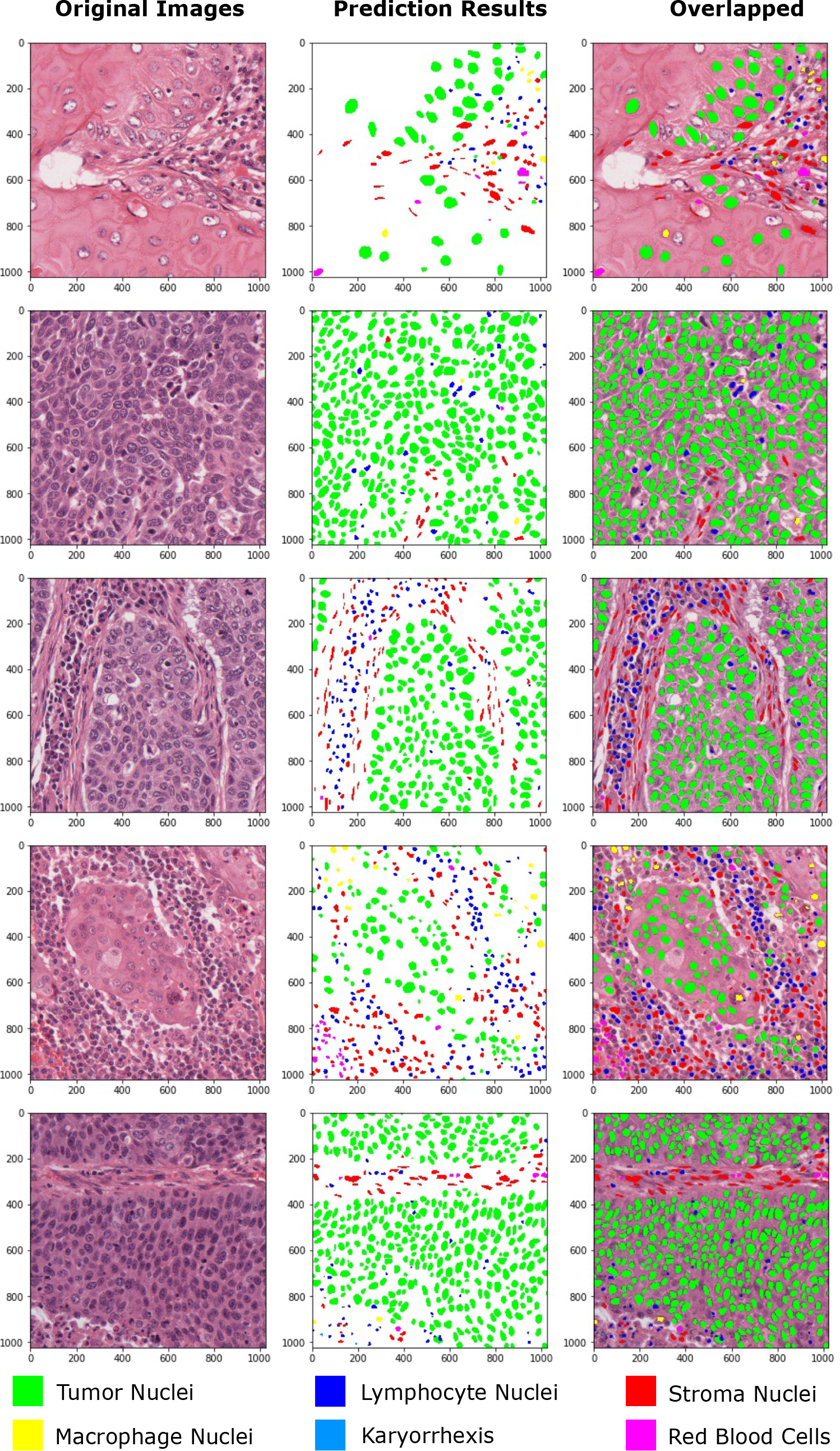
Deep learning based nuclei segmentation results in TCGA lung squamous cell carcinoma pathology images.

